# Evidence and role for bacterial mucin degradation in cystic fibrosis airway disease

**DOI:** 10.1101/047670

**Authors:** Jeffrey M. Flynn, David Niccum, Jordan M. Dunitz, Ryan C. Hunter

**Affiliations:** Department of Microbiology & Immunology, University of Minnesota, 689 23^rd^ Avenue SE, Minneapolis, MN 55455; Division of Pulmonary, Allergy, Critical Care & Sleep Medicine, University of Minnesota, 420 Delaware St. SE, Minneapolis, MN 55455

## Abstract

Chronic respiratory infections are composed of complex microbial communities that incite persistent inflammation and airway damage. Despite the high density of bacteria that colonize the airways, nutrient sources that sustain bacterial growth *in vivo* are unknown. Here we examine the role of respiratory mucins in the ecological dynamics of the cystic fibrosis lung microbiota. While *P. aeruginosa* was unable to efficiently utilize mucins, saliva-derived anaerobes stimulated the growth of opportunistic pathogens when provided mucins as the sole carbon source. The fermentative metabolisms of these oral anaerobes generated amino acids and short chain fatty acids (propionate and acetate) during mucin enrichment *in vitro,* which were also found within expectorated sputum from CF patients. The significance of these findings was supported by *in vivo P. aeruginosa* gene expression, which revealed a heightened response to propionate. Given that propionate is exclusively derived from bacterial fermentation, these data support a central role for mucin fermentation in the carbon flux of the lower airways. More specifically, commensal oral bacteria may contribute to airway disease by degrading mucins, in turn providing nutrients for pathogens otherwise unable to obtain carbon in the lung.

---

Culture-independent surveys of the lung microbiome have revealed a far more complex bacterial community than previously appreciated^1,2,3^. Both in health and disease, the airways harbor a diverse microbiota comprised of multiple taxa infrequently detected by clinical lab culture. While the temporal dynamics of these communities and their association with disease states have been studied in detail, the *in vivo* host environment, and microbial metabolisms therein, is relatively understudied. For example, the means by which lung pathogens obtain sufficient growth energy is not known. The accumulation of mucus secretions in the airways of cystic fibrosis patients represent a potentially abundant bioavailable nutrient source; the major macromolecular constituents, mucins, are a large reservoir of both carbon and nitrogen, and have been measured at concentrations of up to 10 g/L in sputum^4^. These high molecular weight glycoproteins are highly recalcitrant to bacterial degradation^5^, yet, they can serve as a major nutrient source for niche-specific microbiota of the gut and oral cavity. For example, oral streptococci produce a variety of glycolytic and proteolytic enzymes that liberate bioavailable carbohydrates from salivary mucins^6^. In turn, these primary degraders modify the metabolic landscape of the oral cavity and stimulate the growth of secondary colonizers^7^. Similar interactions between commensal gut microbiota are also known^8^. By comparison, the role of airway mucins as a nutrient source for opportunistic pathogens is poorly understood. Here, we investigated mucins as a primary airway carbon source and characterized their potential to sustain pathogen growth. Our results demonstrate that the primary pathogen of CF, *Pseudomonas aeruginosa*, is unable to use mucins directly. This lack of growth raised the question: can other members of the CF microbiota degrade mucins and alter the nutritional reservoir available for pathogen growth? We subsequently found that co-culture of pathogens with oral-derived anaerobes can stimulate pathogen growth using mucins as a sole carbon source. Finally, we revealed that *P. aeruginosa* expresses pathways for the catabolism of mucin-derived fermentation byproducts *in vivo*, strongly supporting a pathogenic role for commensal anaerobes in the progression of CF lung disease.

## Pseudomonas aeruginosa utilizes mucin inefficiently as a sole carbon source

We first tested whether mucins alone could sustain the growth of *P. aeruginosa*. To do so, we assayed strain PA14 for growth in a defined medium containing intact mucins from porcine gastric mucin (PGM) as the sole carbon source (Figure 1a). Interestingly, 10g/L mucin (sugar equivalent of ~45mM glucose assuming 80% sugar by weight) only resulted in a moderate OD_600_ gain of 0.2 after 20 h. By contrast, PA14 grew to 0.8 OD on glucose (15mM), which underscores the inability of PA14 to efficiently break down and utilize mucin glycoproteins.

**Figure 1.**
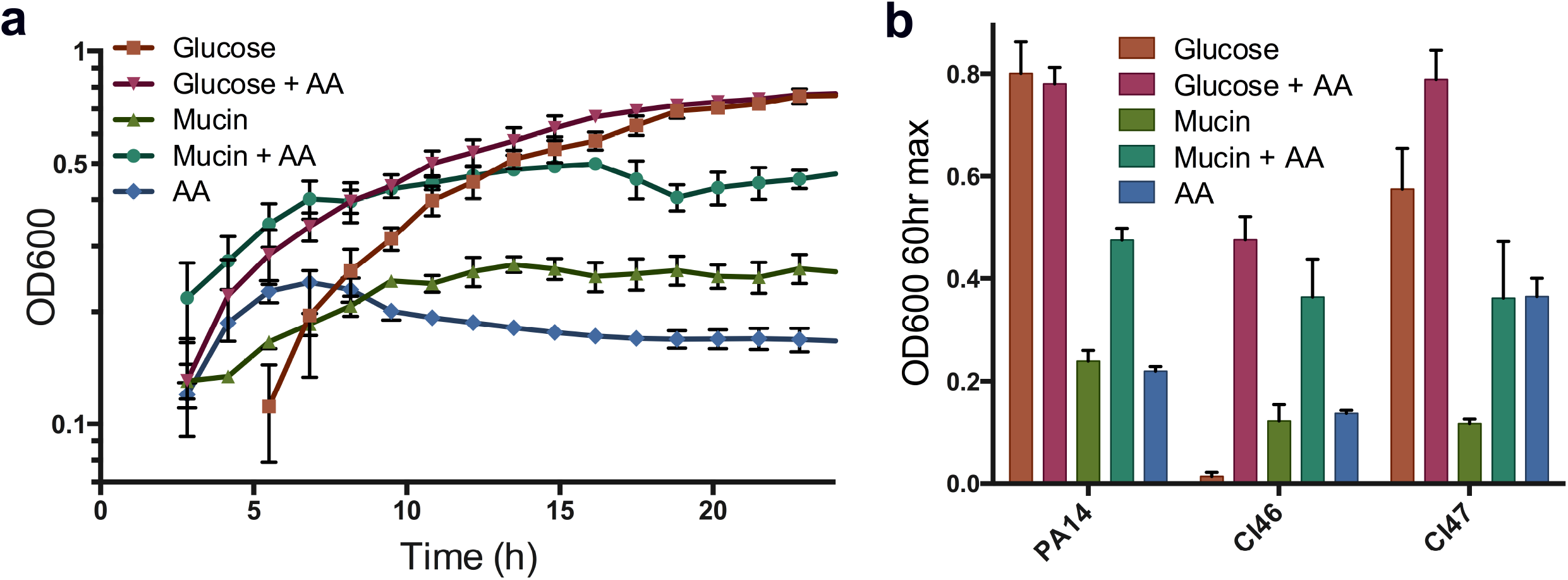
***P. aeruginosa* growth on mucin**. (A) *P. aeruginosa* PA14 grown with mucin and glucose (+/- amino acid supplement). Data are presented as mean values from four independent measurements. (B) *P. aeruginosa* clinical isolates (CI46 and CI47) and PA14 grown with mucin and glucose (+/- complete amino acids supplement) for 40hr. Data are shown as mean +/- SEM of endpoint change in optical density (600nm)(n=4).

Given that the PA14 laboratory strain was isolated from a burn wound^9^, we reasoned that lung-derived clinical isolates would show enhanced growth on mucins. Growth yield of clinical isolates was assayed long past the onset of stationary phase (60 h) to ensure there was no long lag-phase or slow growth phenotypes. Under these conditions, CF-derived *P. aeruginosa* strains CI46 and CI47 grew to an even lower density on mucin relative to PA14 (Figure 1b). Growth on mucin was also compared to growth with glucose alone, with and without the addition of an amino acid supplement to account for auxotrophy, a common trait among CF isolates^10^. CI47 grew well with and without amino acids while CI46 grew poorly on glucose alone, suggesting an auxotrophy that was corrected by the supplement. The ability of the clinical isolates to grow on glucose supplemented with amino acids to a greater extent than the more nutrient-dense mucin medium suggested that the lack of robust growth on mucin was not due to a slow-growth phenotype. Rather, these data suggest that *P. aeruginosa*, including isolates derived from the mucin-rich CF lung environment, are unable to efficiently utilize these complex glycoproteins in monoculture.

## Saliva-derived mucin fermenting bacteria support the growth of opportunistic pathogens *in vitro*

The inefficient use of mucins by *P. aeruginosa* motivated us to consider recent culture-independent studies of the lung microbiota for insights into the carbon flux of the airways. Notably, obligately anaerobic taxa associated with the oral cavity (*e.g. Prevotella, Veillonella)* have gained attention for their abundance in both healthy and diseased lungs^11-13^. A role for these organisms in CF disease is controversial^14,15^; however, they have been characterized for their ability to degrade and ferment both salivary and intestinal mucins^16,17^, in turn supporting the growth of secondary colonizers. Based on these observations, we hypothesized that oral-associated taxa alter the metabolic landscape of the airways by degrading respiratory mucins. More specifically, we predicted that the fermentative degradation of mucin glycoproteins liberates sugars, amino acids and short chain fatty acids (SCFAs) that are readily utilizable by *P. aeruginosa* and other lung pathogens^18^.

To test whether fermenting taxa could stimulate CF pathogen growth, a saliva-derived bacterial community was first enriched on mucin (Supplementary Figure 1). The enrichment culture was then used to inoculate an anaerobic minimal mucin medium supplemented with 1.0% agar to mimic a high viscosity sputum gel^19^ (Figure 2a). Once solidified, PA14 was resuspended in phosphate-buffered saline agar without mucin, and was overlaid on top of the mucin-enriched bacterial community. Under these conditions, PA14 should only grow if provided with diffusible metabolites from the lower phase. After 60 h growth, turbidity was noticeable in the lower phase and a yellow-green pigment characteristic of *P. aeruginosa* was observed throughout the co-culture tube (Figure 2A). By contrast, in the absence of oral anaerobes, little pigment was produced. Colony counts were also determined for *P. aeruginosa, Achromobacter xylosoxidans*, and *Staphylococcus aureus* grown in the presence and absence of mucin-enriched bacteria (Figure 2b). For each pathogen, significant differences (>10-fold; *p*<0.01) in growth were observed in the presence of mucin fermenters relative to tubes in which oral anaerobes were omitted. These data support the hypothesis that oral-derived microbiota can serve as primary mucin degrading organisms in the CF airways, in turn liberating metabolites that can support the growth of opportunistic pathogens.

**Figure 2.**
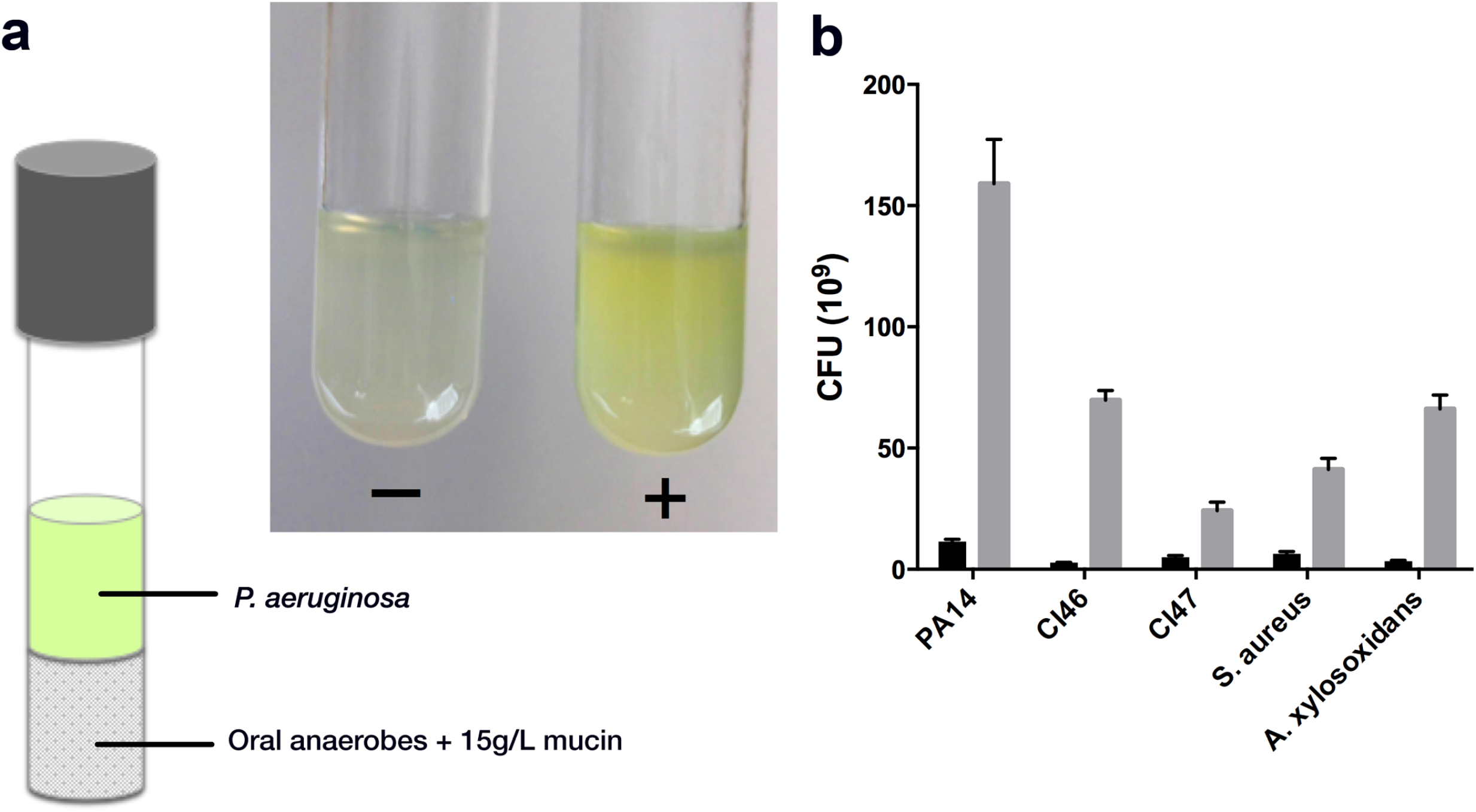
**Oral-derived mucin fermenters support pathogen growth using mucin as the sole carbon source**. (A) Saliva-derived bacteria were grown anaerobically in 1% agar with mucins and overlaid with an immobilized *P. aeruginosa* culture in an agar gel without mucins. After 60hr of growth, co-cultures (inset) containing mucin fermenting anaerobes (+) were pigmented, whereas tubes lacking the fermenting innoculum (-) were not. Pigment is indicative of *P. aeruginosa* growth. (B) Co-culture tubes containing PA14, CI46, CI47, *S. aureus*, and *A. xylosoxidans* overlays in the presence (□) and absence (□) of mucin fermenters. All strains showed significant (*p*<0.01) growth differences when cultured with a fermenting community. Data are presented as mean values +/- SEM from three biological replicates.

## Oral-associated bacteria are present in sputum and are enriched in anaerobic mucin fermentation cultures

To determine if there exists a fraction of the CF lung microbiota that has the ability to degrade and ferment mucins, we performed enrichment experiments on sputum from 14 stable, non-exacerbating CF patients. To do so, a small fraction of sputum was used to inoculate an anaerobic culture with mucins supplied as the sole carbon source. Following growth, genomic DNA was isolated from the initial sputum samples and corresponding enrichment cultures followed by 16S rRNA gene sequencing to identify mucin-fermenting taxa. Consistent with previous studies, sputum (prior to enrichment) harbored a diverse microbiota (Figure 3a) that was highly variable between patients. As expected, *Pseudomonas sp*. made up the highest percentage of sequence reads with a per-patient average of 31.3% across the cohort. Notably, taxa previously characterized for their mucin-degrading activity (*Prevotella, Veillonella, Streptococcus*, and *Fusobacterium)* also made up 35.1% of the population (11.4%, 9.7%, 9.9%, and 4.1%, respectively).

**Figure 3.**
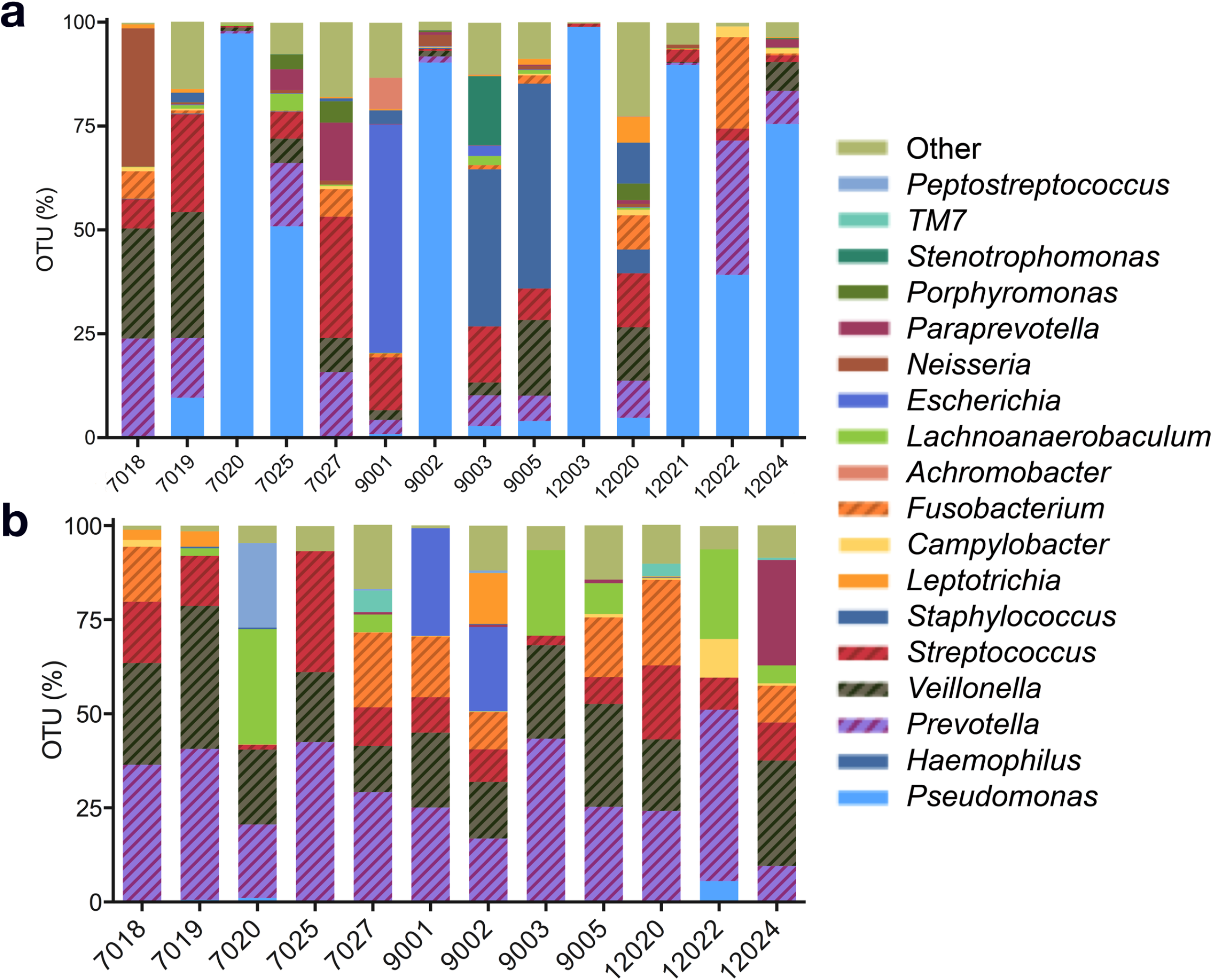
**Mucin enrichment of sputum-derived bacterial communities**. Taxonomic analysis of bacterial communities in (a) sputum and (b) mucin enrichment cultures. A total of 12 enrichment cultures were obtained from 14 cultures seeded with sputum; #12003 and #12021 did not grow in enrichment conditions, accounting for the disparity between sample numbers Oral-associated anaerobes (cross-hatching) capable of mucin fermentation are significantly enriched (*p*<0.05) during growth on mucins under anoxic conditions. Data shown are genus-level abundances.

Post-enrichment, communities were predominated by oral-associated fermenting organisms (Figure 3b). On average, enrichment communities were composed of 73% of taxa known to have mucin degradation ability (27.4% *Prevotella*, 19.2% *Veillonella*, 10.7% *Streptococcus*, 8.4% *Fusobacterium*) with all other genera present at 4% or below. *Lachnoanaerobaculum* and *Prevotella* were significantly enriched on mucins while *Neisseria, Staphylococcus*, and *Pseudomonas* were selected against (*p*<0.05). The high composition of mucin-fermenting taxa in the initial sputum samples indicates a suitable niche space exists in the CF lung for oral-associated mucin fermenting anaerobes. The enrichment results demonstrate that these taxa found within sputum retain their mucin fermentation capacity during growth in the lower airways, and could potentially cross-feed lung pathogens *in vivo*.

## Mucin fermenting taxa are present within explanted CF lungs

The presence of oral flora in the lower airways and their contribution to CF lung disease progression is controversial due to the possibility of contamination from the route of sampling^14,15^. To address this concern, we sampled mucus directly from multiple lobes of three explanted pairs of lungs derived from late-stage CF patients (Supplementary Table 1). Not surprisingly, each lobe from all three patients harbored an abundance of *Pseudomonas* sp., which is characteristic of late-stage lung disease^14,20,21^. Yet, two patients also contained a notable proportion of mucin-fermenting taxa identified in our enrichment experiments. For example, the left lingula (left middle lung) from patient A contained relatively diverse microbiota, of which approximately 4.7% were *Prevotella, Streptococcus*, and *Veillonella*. Patient B harbored known mucin-fermenting taxa (~7.9% of sequence reads) in the lower left lung. Patient C was dominated (>99%) by *Pseudomonas*, but even in this patient, it is noteworthy that mucin-fermenting anaerobes were associated with several regions of the airways. Together, these data support that a proportion of oral-associated anaerobes in our sputum samples were derived from the lower airways.

## Sputum-derived fermenting organisms produce short-chain fatty acids and amino acids when grown on mucin

To study the cross-feeding ability of mucin fermenters in further detail, we characterized the metabolites that were generated during anaerobic mucin enrichment. To do so, we used targeted high performance liquid chromatography (HPLC) for the quantification of organic acids produced in mixed acid fermentation (acetate, butyrate, citrate, formate, isobutyrate, isovalerate, ketobutyrate, ketoisovalerate, lactate, 2-methylbutyrate, propionate, succinate and valerate), and gas chromatography for the quantification of amino acids. Mucin-enriched sputum cultures generated high amounts of acetate (30.2 +/- 9.7 mM) and propionate (15.4 +/- 3.8 mM), while only one enrichment sample containing an abundance of *Streptococcus sp*. (#7025, see Figure 3a) produced a detectable concentration of lactate (7.8 mM) (Figure 4a). No other SCFAs assayed were found in the mucin-enrichment supernatants. Not surprisingly, free amino acids were also present in each sample (1.37 +/- 1.32 mM total) (Figure 4b), suggestive of their liberation from the mucin polypeptide backbone. The abundance of amino acids correlates well with previous studies of sputum composition where they were found to be present in appreciable quantities^22^. These data demonstrate that acetate, propionate, and amino acids are byproducts of mucin fermentation by sputum-derived anaerobes and suggest that they may be bioavailable carbon sources that can cross-feed pathogens *in vivo*.

**Figure 4.**
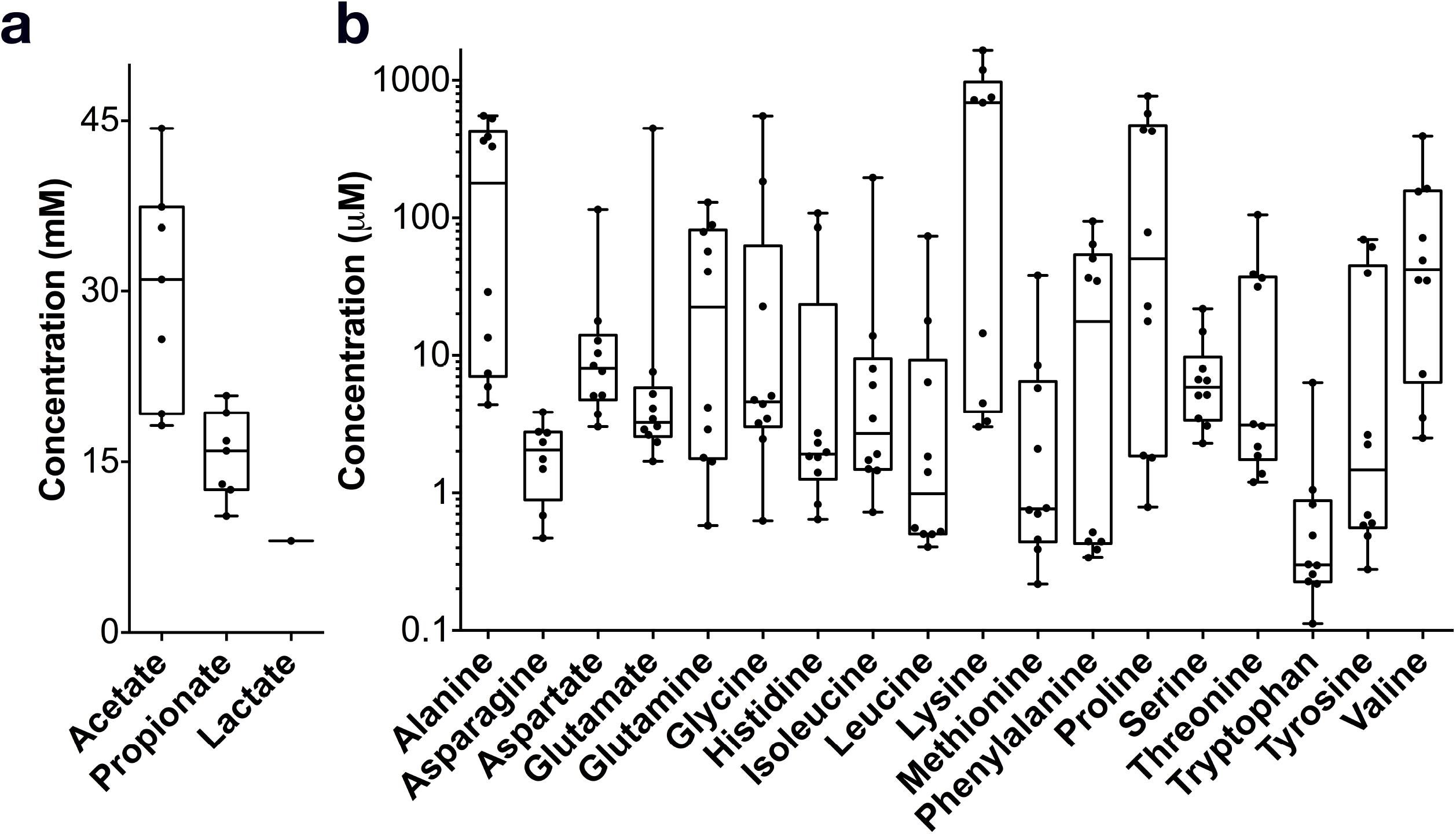
**Mucin-enrichment metabolite profiling**. (a) short chain fatty acids (n=8) and (b) amino acids (n=10) were quantified in mucin enrichment cultures. Arginine and cysteine were undetectable due to their co-elution with other metabolites.

## Cystic fibrosis sputum contains short-chain fatty acids and *Pseudomonas aeruginosa* responds to propionate *in vivo*

16S rRNA gene analyses demonstrated the presence of a fermentative bacterial community in the CF airways. To provide further evidence of fermentative activity *in vivo*, we used two complementary techniques: mass spectrometry and quantitative reverse transcription PCR (qRTPCR). First, using gas chromatography mass spectrometry (GC-MS), acetate and propionate were quantified in paired saliva/sputum samples collected from 9 stable CF patients. These data revealed no significant differences for either SCFA between the oral cavity and lung (Supplementary Figure 2). Given the presence of fermenting taxa and SCFA previously identified in saliva^23^, this result was not unexpected. To circumvent the effect of SCFA contaminants from the mouth during sampling, an oral rinse was first performed, prior to saliva and immediate sputum collection (see Methods). Using this sampling strategy on 7 additional patients, acetate concentrations were significantly higher in sputum relative to saliva (5.9 +/- 1.8 mM versus 2.4 +/- 1.8, *p*<0.01)(Figure 5a), consistent with metabolites generated during enrichment culture (see Figure 4A) and indicative of a fermentative bacterial community within the CF lung. Propionate, on the other hand, showed no significant difference (*p*=0.89) between paired samples (Figure 5b). This result was surprising given the high levels of propionate generated during in vitro mucin enrichments. Because of this disparity, we hypothesized that propionate, over acetate, is preferentially utilized by members of the CF lung community in a cross-feeding relationship.

**Figure 5.**
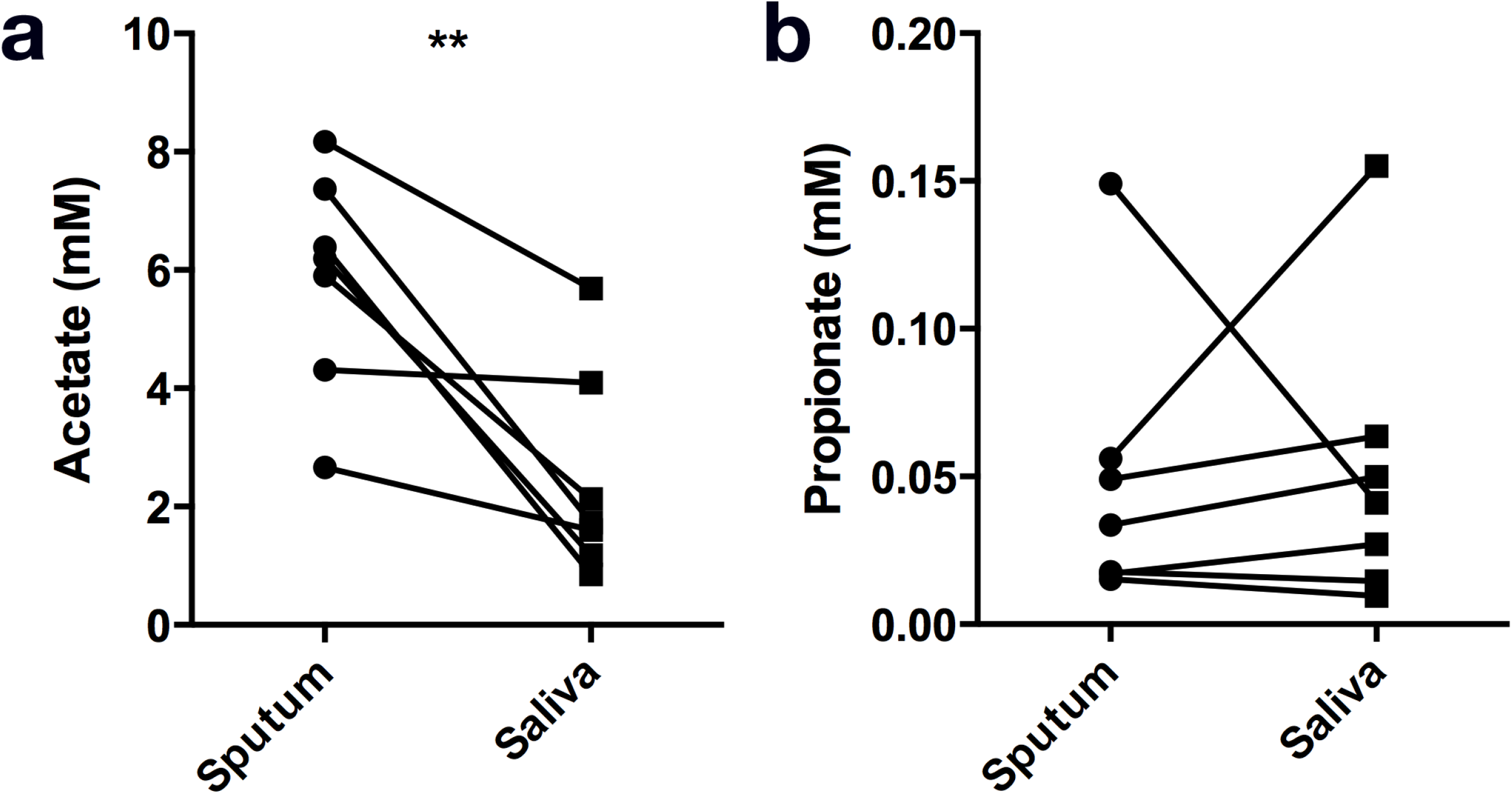
**Evidence for mucin-derived short chain fatty acid production *in vivo.*** Direct measurements of (a) acetate and (b) propionate in paired sputum and saliva samples from CF patients. Acetate concentrations were significantly higher in sputum (*p* = 0.006, **).

If SCFAs were simultaneously produced and consumed by the airway bacterial community, we would expect the concentration of these metabolites to remain low, and genes required for their catabolism to be highly expressed. To address this possibility, we used qRTPCR to measure *in vivo* bacterial gene expression. Genomic content suggests that *Pseudomonas* has committed pathways for both acetate and propionate catabolism (encoded by *acsA/B* and *prpBCD*, respectively). Therefore, as a proxy of the use of mucin-derived metabolites by *P. aeruginosa*, we targeted the *in vivo* expression of *acsA* (encoding acetyl-coA synthetase) and prpD (methylcitrate lyase) relative to their expression levels under controlled conditions. In vitro, *acsA* and *prpD* were differentially expressed by PA14 in the presence of acetate (5.8-fold higher, *p*=0.02) and propionate (11.4-fold, *p*<0.01), respectively, relative to growth on glucose alone (Figure 6). When both SCFAs were present, prpD was expressed significantly higher than *acsA* (*p*<0.01), which were both upregulated relative to the glucose control (2.4 and 4.8-fold for *acsA* and *prpD*, respectively). When compared to in vitro cultures, analysis of sputum revealed that prpD (5.4-fold, *p*=0.02) but not *acsA* (no change, *p*=0.98) was upregulated. These data are indicative that propionate is generated in the airways of CF patients and that *P. aeruginosa* both senses and catabolizes propionate *in vivo*, providing further evidence that mucin-associated fermentative metabolisms contribute to the metabolic landscape of the CF lung.

**Figure 6.**
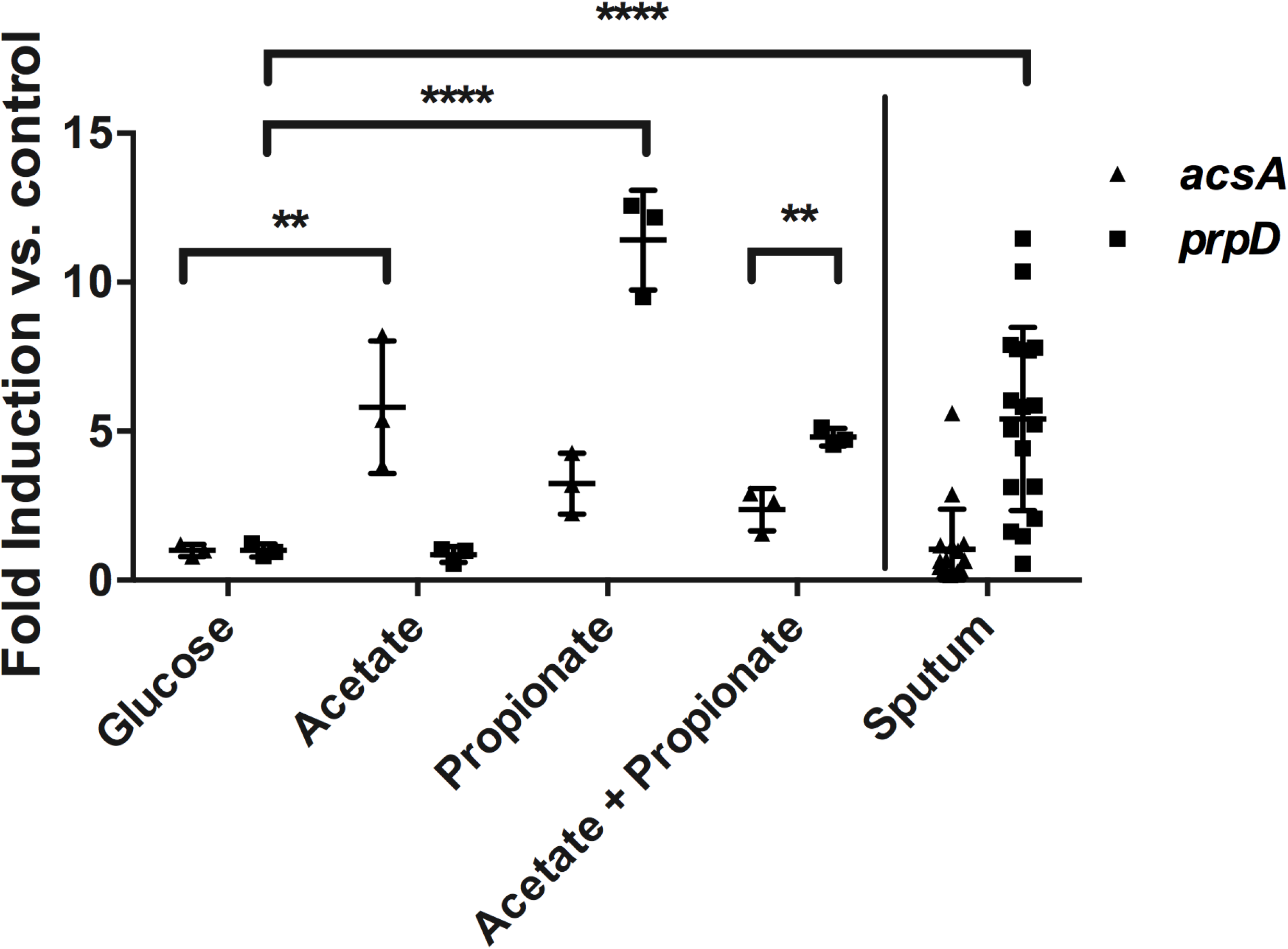
*In vitro* expression (left) of *acsA* (▴) and *prpD* (■) are upregulated in response to acetate and propionate, respectively. *prpD* is preferentially expressed in the presence of both acetate and propionate. Data shown are biological triplicate measurements for PA14. *In vivo* (right), *prpD* is upregulated (5.4 fold ± 3.1) while *acsA* is not (1.0 fold ± 1.3 SD) (*n*=17). *p* ≤ 0.01, **; *p* ≤ 0.001, ***; *p* ≤ 0.0001, ****.

## DISCUSSION

Metabolite constituents of expectorated sputum are known to support bacterial growth *in* vitro^22,24,25^ but the nutritional role of intact mucins has not been characterized. Previous studies reported that mucins support the carbon demands of *P. aeruginosa in vitro*^26,27^; however, these studies included autoclaved preparations of commercial PGM that contain low molecular mass impurities. In fact, when PGM was filtered and dialyzed in our study (leaving only intact, large glycoproteins), *P. aeruginosa* inefficiently utilized mucins as a carbon source on their own. Rather, in the presence of purified mucin, fermentative oral-associated bacteria were required to stimulate pathogen growth. Moreover, we revealed that *in vitro* mucin fermentation generated high concentrations of SCFAs and amino acids, which were both abundant (acetate) and utilized by *P. aeruginosa* (propionate) in patient sputum. Together, these data provide strong evidence for a central role of mucin fermentation in the carbon flux of lower airway disease and define a pathogenic role for oral-associated anaerobes in the progression of CF-associated airway disease.

*In vitro*, degradation of O-linked glycans and the polypeptide backbone of mucin by sputum-derived anaerobes generated an abundance of SCFAs and amino acids. Given the known substrate utilization profile of *P. aeruginosa*^18^, these compounds also likely sustained pathogen growth within the co-culture experiment. Consistent with recent studies^32,33^, SCFAs were also found in sputum across our CF patient cohort. Because SCFAs serve as a reliable biomarker of fermentative metabolism^34^, their universal presence suggests that bacterial fermentation is a ubiquitous metabolic activity in the CF airways. More specifically, our data suggest that the fermentation of mucins is a likely origin for metabolites that have been shown to support growth of *Pseudomonas*^22^ in expectorated sputum. Moreover, the high levels of *prpD* expression confirms that *P. aeruginosa* is both sensing and utilizing propionate *in vivo*, suggesting that the lower-than-expected concentrations of this metabolite in sputum can be explained by its consumption as opposed to a lack of production within the airway environment.

Our data create a compelling model for the role of oral anaerobes in the development of CF lung disease, whereby pathogens cannot become established until mucin fermenters have colonized the lower airways. In CF, impaired mucociliary clearance and defective immunological responses increase the likelihood of oral anaerobe colonization and the degradation of respiratory mucins. In turn, anaerobes left uninhibited can condition the lung environment into a niche that is suitable for pathogens to thrive. Several lines of clinical evidence support this model: (i) pediatric patients have asymptomatic primary colonization by oral anaerobes^35,36^ prior to the establishment of chronic *P. aeruginosa* infections, (ii) routine administration of broad-spectrum antibiotics is an effective therapy to delay the onset of chronic colonization by *P. aeruginosa* and reduce the frequency of acute exacerbations^37,38^, and (iii) the *in vitro* antibiotic susceptibility of lung pathogens does not correlate with clinical outcomes^39^. In the latter instance, we suspect that antibiotics do not solely target the pathogen, but rather disrupt the complex metabolic interactions that supply them with substrates for growth and stimulate their pathogenicity.

This work raises the question of the importance of mucin-degrading anaerobes in late stages of CF disease when bacterial diversity often declines and becomes predominated by *P. aeruginosa*^40^. In severe disease states, it is known that chronic airway infections can lead to a ‘leaky’ lung epithelium that allows bioavailable metabolites to reach the epithelial surface^41^. Additionally, persistent *P. aeruginosa* infections incite a neutrophil-dominant inflammatory response that is associated with increased concentrations of host-derived proteases^42^. Human neutrophil elastase, for example, is capable of cleaving glycoproteins into free amino acids, and has been implicated in the progression of lung disease^43^. Moreover, neutrophilic burst may also contribute to the bioavailable nutrient pool, potentially diminishing the need for anaerobic fermenters as *P. aeruginosa* populations colonize the airway niche and incite a heightened immune response. We are currently investigating the temporal dynamics of both host and microbiota mucin degradation across the spectrum of disease severity.

Altogether, this study underscores the importance of defining the underlying microbial ecological dynamics associated with airway disease. In particular, our data warrant further studies of targeted therapies towards fermentative organisms and their metabolisms that may contribute to the establishment and progression of chronic CF lung infections. In a broader context, the presence of both oral anaerobes and known pathogens in other, mucus rich environments – chronic obstructive pulmonary disease, chronic sinusitis, ventilator-associated pneumonias – suggests that mucin fermentation and metabolic cross-feeding may be a widespread disease phenomenon. A more detailed understanding of these metabolic interactions are vital to the development of effective therapeutic strategies and improved clinical outcomes.

## Methods

### Bacterial strains and culture conditions

*P. aeruginosa* PA14 and *Staphylococcus aureus* MN8 were obtained from D.K. Newman (California Institute of Technology). Clinical strains of *P. aeruginosa* (CI46, CI47) and *A. xylosoxidans* MN001^44^ were isolated from patients enrolled in this study. Strains were routinely cultured in Luria Bertani medium or a minimal mucin medium containing 60mM KH_2_PO_4_ (pH 7.4), 90mM NaCl, 1mM MgSO_4_, 10 g L^-1^ (or 15 g L^-1^ where specified) porcine gastric mucin (Sigma), and a trace minerals mix^45^. During preparation, mucin was autoclaved, dialyzed using a 13 kDa molecular weight cutoff membrane, clarified by centrifugation, followed by passage through a 0.45 μm Millipore filter to sterilize soluble intact glycoproteins. Glucose and NH4Cl were supplemented at 13mM and 60mM, respectively, where specified. MEM essential and non-essential amino acid mixes (Sigma) were added at a final concentration of 0.5X the recommended concentration. For enrichment growth, ~100 μL of sputum was used to inoculate minimal mucin medium under anoxia (95% N_2_, 5% CO_2_) and the remainder was frozen at −80°C. Following 48h of incubation at 37°C, 100 μL of culture was used to inoculate a second anaerobic culture tube and allowed to grow for 48h. Genomic DNA was isolated from both the initial sputum sample and enrichment cultures, and bacterial community composition was determined using 16S rRNA gene sequencing.

### Patient cohort and sample collection

Forty-eight participants with CF were recruited at the University of Minnesota Adult CF Center and informed consent was obtained for all subjects. For enrichment cultures, sputum was expectorated into 50ml conical tubes, placed on ice, and processed within three hours. Sputum used for qRTPCR analysis was placed in RNALater (Sigma) immediately following expectoration, and stored at −80°C for further processing. For metabolite analysis (n=9), paired saliva and sputum samples were also collected in conical tubes and immediately stored at −80°C. For a second set of samples, subjects (n=7) first performed an oral rinse, followed by saliva then sputum collection less than 30 seconds afterwards. To obtain clinical isolates of *Pseudomonas sp*., sputum aliquots were cultured on *Pseudomonas* Isolation Agar (Oxoid) for 72 hours at 37°C. Colonies were screened using PCR^46^. Isolates positively identified as *P. aeruginosa* were stored in 15% glycerol and frozen at −80°C.

Lung tissue was obtained from three late-stage CF patients undergoing double lung transplantation. Upon extraction, explanted lung parenchyma was separated from the main bronchi and dissected into six lobes (right upper, right middle, right lower, left upper, left lingula, left lower) by the UMN BioNet Tissue Procurement Facility. Mucus was extracted via syringe from the conducting airways of each lobe, and frozen at −80°C for subsequent 16S rRNA gene profiling. These studies were approved by the Institutional Review Board at the University of Minnesota (UMN IRB nos. 1401M47262 and 1404M49426).

### DNA extraction and 16S community analysis

Lung mucus, sputum and enrichment cultures were thawed to room temperature, and 500 μL of each sample was used for genomic DNA extraction using the PowerSoil DNA isolation kit (MoBio, Carlsbad, CA). Purified DNA was submitted to the University of Minnesota Genomics Center (UMGC) for 16S library preparation using a two-step PCR protocol. Briefly, the primary amplification step was performed using KAPA HiFi Polymerase (Kapa Biosystems, Woburn, MA) and primers specific for the region flanking 16S rRNA V4 region that included adapter tails for adding indices in a secondary amplification step (Supplementary Table 2). Amplicons were diluted 1:100, and a subsequent PCR reaction was performed to add indices and Illumina flow cell adapters. For each sample, a pair of 8 base bar codes were included to distinguish between samples. Indexing primers are listed in Supplementary Table 2. Amplicons were quantified using PicoGreen (Life Technologies, Carlsbad, CA), normalized, pooled, and ~1 μg of material was concentrated to 10μl using 1.8x AMPureXP beads (Beckman Coulter, Brea, CA). The pooled sample was size selected at 427bp +/- 20% on a Caliper XT DNA 750 chip (Caliper Life Science, Hopkinton, MA). Amplicons were cleaned using AMPureXP beads and analyzed using an Agilent Bioanalyzer Chip (Agilent Technologies, Santa Clara, CA). The final pooled sample was then diluted to 2 nM, 10μl was denatured using 0.2N NaOH, diluted to 8pM in HT1 buffer, spiked with 15% phiX, heat denatured for 2 min, and sequenced using a MiSeq 600 cycle v3 kit (Illumina, San Diego, CA). A defined mock community and reagent blanks were also sequenced as controls.

Raw sequence reads were obtained from UMGC and analyzed using a QIIME^47^ pipeline developed by the UMGC. Briefly, read pairs were stitched together and 16S amplicon primers were removed using PandaSeq (version 2.7)^48^. Fastq files were merged and sequence IDs converted to QIIME format using a custom perl script. Chimeric sequences were detected using the QIIME (version 1.8.0) script identify_chimeric_seq.py function, using the usearch61 method. Open reference OUT picking was performed using the pick_open_reference_otus.py script, using the usearch61 method and the Greengenes 13_8 16S rRNA reference database^49^ clustered at 97% similarity. MetaStats was used to detect differentially abundant features of the mucin-enriched bacterial community^50^. All fastq files are available online at the NIH Sequence Read Archive (http://www.ncbi.nlm.nih.gov/bioproject) under BioProject accession number SRP067035.

### *In vitro* cross-feeding

A saliva-derived mucin-fermenting bacterial community (Supplementary Figure 1) was used to inoculate 2 mL of mucin (15g/L) minimal medium in molten 1% agar at 50ºC in a glass culture tube under anoxic conditions (Fig. 2A). Upon solidification, an additional 2 mL of molten minimal medium agar (without mucin) was inoculated with a 1/10000 dilution of an overnight culture of *P. aeruginosa* (or other strains where specified). Samples were then poured over the mucin fermenting community. Tubes without mucin-fermenters were used as controls. After solidification, co-cultures were placed at 37ºC for 65 hours. Agar plugs were then removed from the upper phase, sectioned and normalized by weight, followed by homogenization by pipetting in 1 mL of phosphate buffered saline. Colony forming units were determined by serial dilution and plating on LB agar.

### Metabolite quantification

Targeted quantification of mucin fermentation byproducts (acetate, butyrate, citrate, formate, isobutyrate, isovalerate, ketobutyrate, ketoisovalerate, lactate, 2-methylbutyrate, propionate, succinate and valerate) was performed via high performance liquid chromatography (GC-MS). The system consisted of a Shimadzu SCL-10A system controller, LC-10AT liquid chromatograph, SIL-10AF autoinjector, SPD-10A UV-Vis detector, and CTO-10A column oven. Separation of compounds was performed with an Aminex HPX-87H guard column and an HPX-87H cation-exchange column (Bio-Rad [Hercules, CA]). The mobile phase consisted of 0.05 N H2SO4, set at a flow rate of 0.5 ml min-1. The column was maintained at 50°C and the injection volume was 50 μl. Amino acid and metabolite (acetate and propionate) quantification from enrichment supernatants were performed by Millis Scientific, Inc. (Baltimore, MD) using liquid chromatography-mass spectrometry and gas chromatography-mass spectrometry (GC-MS). Samples for amino acid quantification were spiked with 1 μl of uniformly labeled amino acids (Cambridge Isotope Labs) and derivatized using AccQ-Tag reagent (Waters Corp.) for 10 min at 50°C. A Waters Micromass Quatro LC-MS interfaced with a Waters Atlantis dC18 (3 μm 2.1×100 mm) column was used. Reverse-phase LC was used for separation (mobile phases A:10mM ammonium formate in 0.5% formic acid, B:methanol) with a constant flow rate (0.2 ml/min) and a column temperature of 40°C. Electrospray ionization was used to generate ions in the positive mode and multiple reaction monitoring was used to quantify amino acids. Samples (~100 μl) for acetate and propionate quantification were first diluted (150 μl water), spiked with internal standards [^13^C_2_] of 1000ppm acetate [^13^C_2_] and 1000 ppm propionate [^13^C_1_]) and acidified using 2 μl of 12N HCl. After equilibration at 60°C for 2h, carboxen/ polydimethylsiloxane solid phase microextraction (SPME) fiber was used to adsorb the headspace at 60°C for 30min. Acids were then desorbed into the gas chromatograph inlet for 2 min. A 30 m × 0.32 mm ID DB-624 column attached to a Thermo Electron Trace gas chromatograph with helium carrier gas (2.0 ml/min) was used for separation of analytes. A Waters Micromass Quatro GC mass spectrometer was used for detection and quantification of target ions. Significance between sputum and saliva samples were determined by paired Student’s t-test.

### RNA isolation, purification, and qRTPCR

To quantify *P. aeruginosa* gene expression *in vivo*, sputum (n=17) was expectorated into RNAlater and immediately frozen to preserve the gene expression profile. Frozen sputum was thawed in TriZol (Life Technologies), homogenized using ceramic beads and purified according to the manufacturer’s protocol. RNA was concentrated using the Clean & Concentrator kit (Zymo) and de-salted using Turbo DNA-free (Life Technologies). Bacterial RNA was enriched using the MicrobEnrich kit (Life Technologies), and purity was confirmed using Qubit (LifeTechnologies) spectrophotometry. qRTPCR was performed as previously described^51^. Briefly, DNA was reverse transcribed from 1 μg of total RNA using the iScript cDNA synthesis kit (BioRad). cDNA was then used a template for quantitative PCR on an iQ5 thermocycler (BioRad) using iTaq Universal SYBR Green Super Mix (BioRad). Triplicate measurements were made on each sputum sample. For control cultures, *P. aeruginosa* was grown in 4-morpholinepropanesulfonic acid (MOPS) minimal medium to an OD600 of~0.6 supplemented with glucose (12mM), acetate (20mM), propionate (20mM), or acetate + propionate (20mM each) where specified. After growth, cells were harvested by centrifugation, frozen at −80°C, and RNA was extracted as described above. Primer pairs are listed in Supplementary Table 2. For all primer sets, the following cycling parameters were used: 94°C for 3 min followed by 40 cycles of 94°C for 60s, 55°C for 45s, and 72°C for 60s. Primer efficiencies were tested for *clpX* (91.6%), *acsA* (91.6%) and *prpD* (96.8%). Relative RNA values were calculated from the Ct values reported and the experimental primer efficiencies, and were normalized to the expression of *clpX. clpX* was compared to *oprI* values to ensure constitutive expression levels. Significance between treatments was determined by two-tailed unpaired Student’s t-test.

## Acknowledgements

This work was supported a Pathway to Independence Award from the National Heart Lung & Blood Institute to R.C.H. (R00HL114862), and the National Center for Advancing Translational Sciences (UL1TR000114). J.M.F. is supported by a NIH Lung Sciences T32 fellowship (#2T32HL007741-21). The content is solely the responsibility of the authors and does not necessarily represent the official views of the NIH. We acknowledge the Minnesota Supercomputing Institute (http://www.msi.umn.edu), Sarah Lucas for her computational expertise, Daryl Gohl and John Garbe at the UMN Genomics Center for sequencing assistance, Millis Scientific for mass spec analysis, and Daniel Bond for the use of analytical instruments.

## Author Contributions

J.M.F. performed all experiments except for Illumina library generation and sequencing. D.N. and J.M.D. recruited, consented, and collect samples/tissue from human subjects. R.C.H. and J.M.F. analysed and processed the data. R.C.H. and J.M.F. wrote the manuscript. R.C.H. supervised the project.

---

**Supplementary Figure 1.**
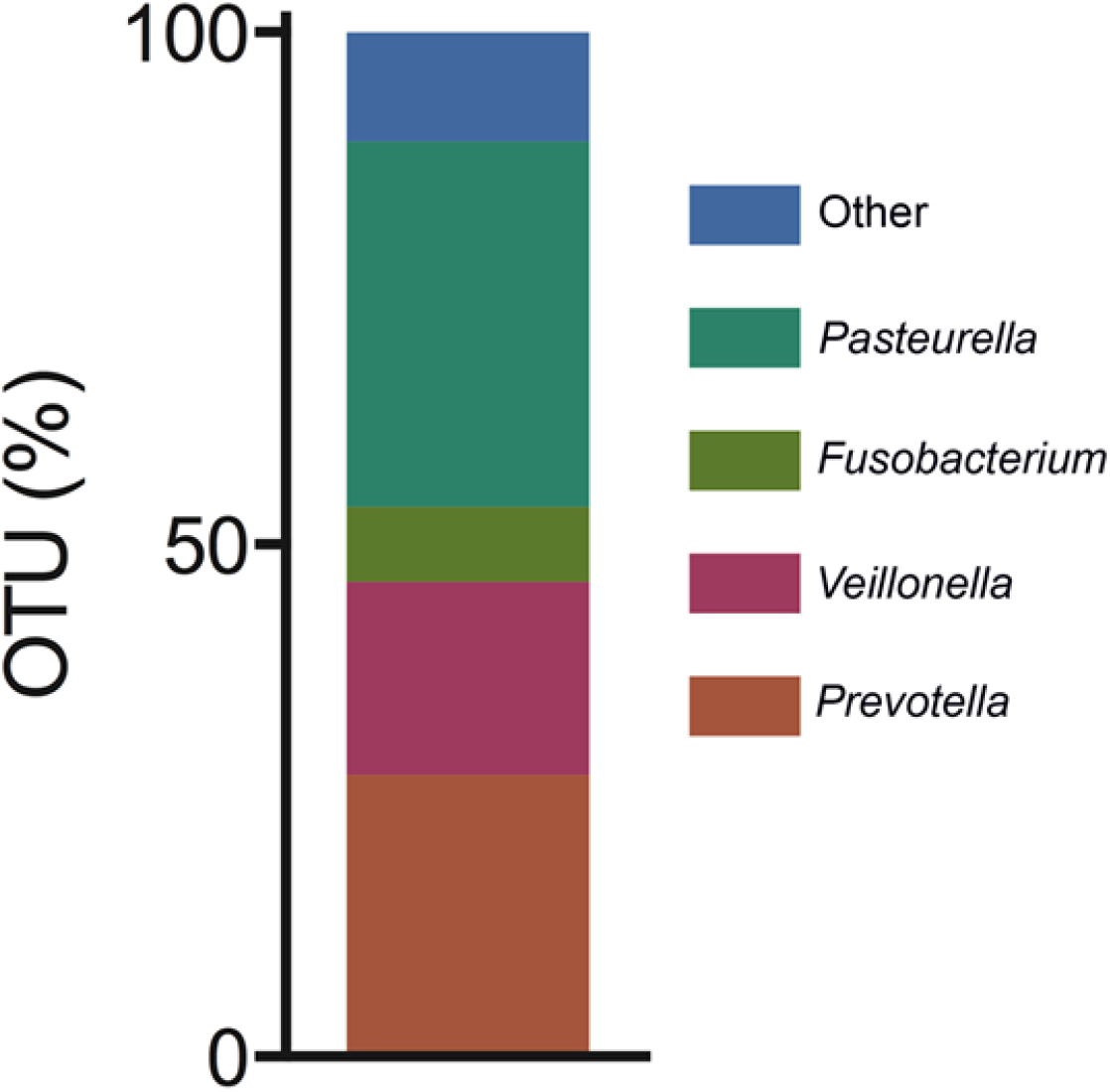
Taxonomic composition of the mucin-enriched, saliva-derived bacterial community used in this study.

**Supplementary Figure 2.**
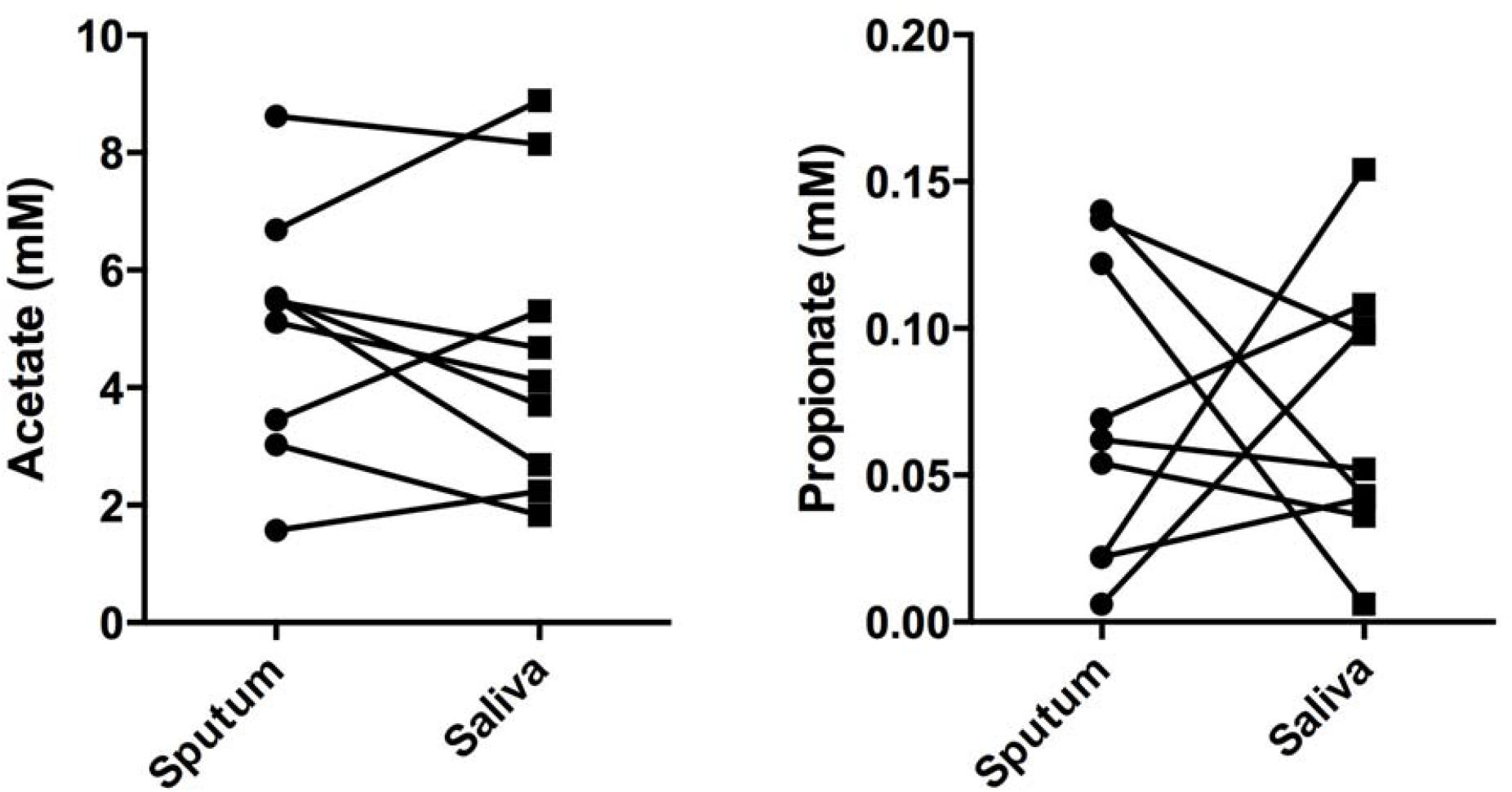
Direct measurements of (a) acetate and (b) propionate in paried sputum and saliva samples between CF patients (n=9) without an oral rinse between sampling sites. There were no significant difference between saliva and sputum for either SCFA (*p*=0.51 and 0.98, respectively).

**Supplementary Table 1.** Relative abundance of bacterial OTUs in explanted CF lungs.

**Patient A**
| Genus<sup>a</sup> | Right Upper | Right Middle | Right Lower | Left Upper | Left Lingula | Left Lower |
| --- | --- | --- | --- | --- | --- | --- |
| *Pseudomonas* | 98.87% | 98.53% | 97.49% | 98.31% | 82.75% | 96.90% |
| *Streptococcus* | 0.01% | 0.05% | 0.14% | 0.09% | 1.96% | 0.03% |
| *Veillonella* | - | 0.04% | 0.10% | - | 0.39% | 0.45% |
| *Prevotella* | - | 0.07% | 0.12% | 0.03% | 2.35% | 0.48% |
| *Haemophilus* | - | - | - | 0.01% | 3.14% | - |
| *Staphylococcus* | 0.02% | 0.04% | 0.06% | 0.08% | 0.20% | 0.15% |
| *Fusobacterium* | - | - | 0.12% | 0.01% | - | - |

**Patient B**
| Genus<sup>a</sup> | Right Upper | Right Middle | Right Lower | Left Upper | Left Lingula | Left Lower |
| --- | --- | --- | --- | --- | --- | --- |
| *Pseudomonas* | 99.75% | 99.32% | 99.26% | 99.88% | 99.34% | 86.46% |
| *Streptococcus* | - | 0.04% | - | 0.02% | - | 1.61% |
| *Veillonella* | - | 0.06% | 0.09% | - | 0.02% | 3.83% |
| *Prevotella* | - | 0.01% | 0.02% | - | 0.01% | 1.45% |
| *Haemophilus* | 0.03% | - | 0.01% | - | - | - |
| *Staphylococcus* | 0.03% | 0.01% | 0.02% | - | 0.04% | 0.31% |
| *Fusobacterium* | - | - | 0.01% | - | - | 0.99% |

**Patient C**
| Genus<sup>a</sup> | Right Upper | Right Middle | Right Lower | Left Upper | Left Lingula | Left Lower |
| --- | --- | --- | --- | --- | --- | --- |
| *Pseudomonas* | 99.89% | 99.87% | 99.88% | 99.82% | 99.86% | 99.86 |
| *Streptococcus* | - | 0.01% | - | 0.01% | - | - |
| *Veillonella* | 0.01% | - | - | 0.05% | - | 0.01% |
| *Prevotella* | 0.01% | - | - | 0.02% | - | - |
| *Haemophilus* | - | - | - | - | - | - |
| *Staphylococcus* | 0.01% | - | - | - | - | - |
| *Fusobacterium* | - | - | - | - | - | - |

<sup>a</sup> Shown are the three most abundant known lung pathogens (*Pseudomonas*, *Haemophilus*, *Staphylococcus*) and the percentage of known mucin-degrading anaerobes (*Prevotella*, *Streptococcus*, *Veillonella* and *Fusobacterium*)

**Supplementary Table 2.**
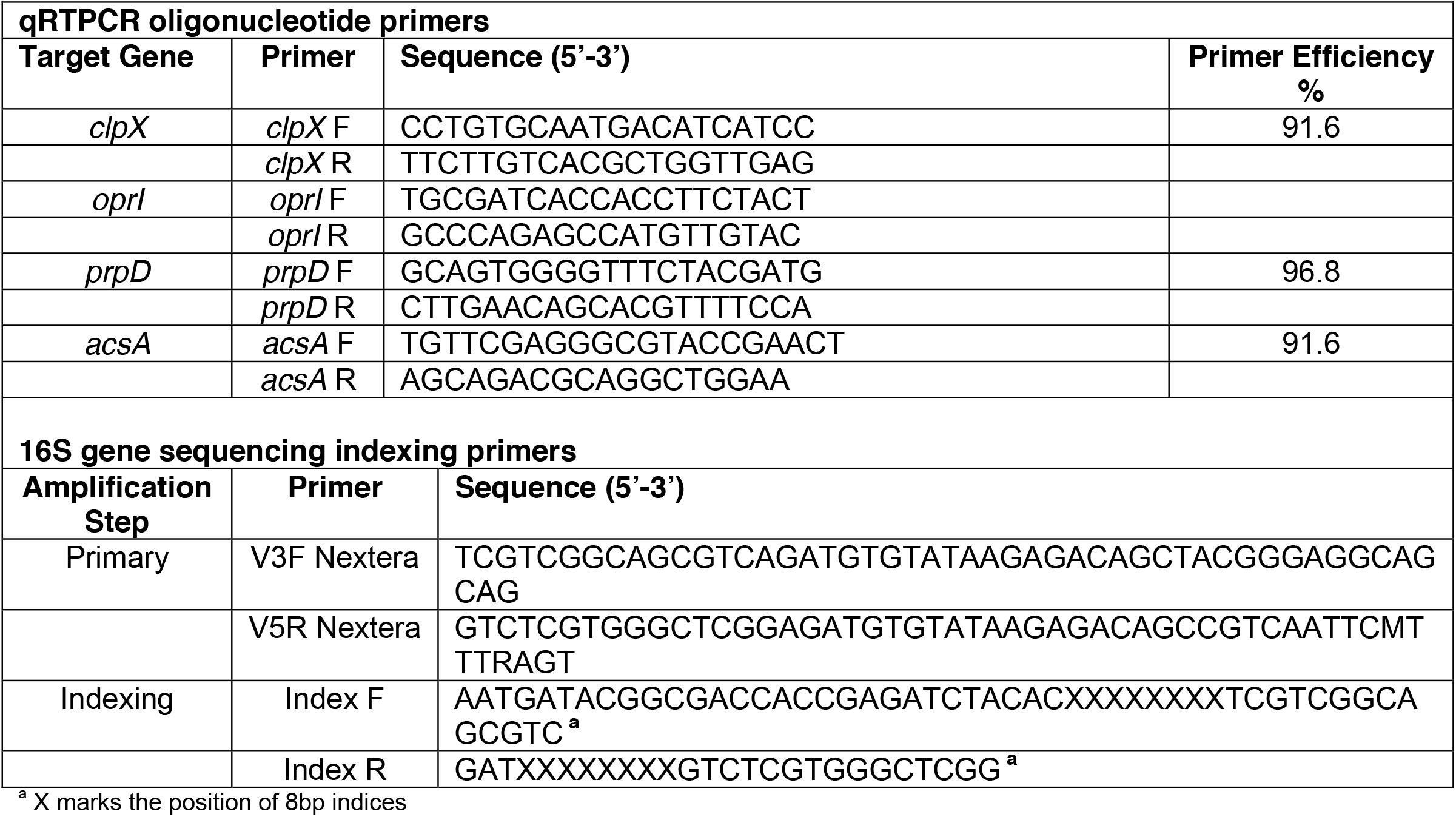
Oligonucleotide primers used in this study.

## References

1 Carmody L. A. et al. Changes in cystic fibrosis airways microbiota at pulmonary exacerbation. Ann. Am. Thorac. Soc. 10, 179–187 (2013).

2 Rogers G. B. et al. Use of 16S rRNA gene profiling by terminal restriction fragment length polymorphism analysis to compare bacterial communities in sputum and mouthwash samples from patients with cystic fibrosis. J. Clin. Microbiol. 44, 2601–2604 (2006).

3 Zemanick E. T. et al. Assessment of airway microbiota and inflammation in cystic fibrosis using multiple sampling methods. Ann. Am. Thorac. Soc. 12, 221–229 (2015).

4 Henderson A. G. et al. Cystic fibrosis airway secretions exhibit mucin hyperconcentration and increased osmotic pressure. J. Clin. Invest. 124, 3047–3060 (2014).

5 Roberton A.M, McKenzie C.G., Sharfe N. & Stubbs L.B. A glycosulphatase that removes sulphate from mucus glycoprotein. Biochem. J. 293, 683–689 (1993).

6 Wickstrom C., Herzberg M.C., Beighton D. & Svensäter G. Proteolytic degradation of human salivary MUC5B by dental biofilms. Microbiol. 155, 2866–2872 (2009).

7 Kolenbrander P. Multispecies communities: interspecies interactions influence growth on saliva as sole nutritional source. Int. J. Oral Sci. 3, 49–54 (2011).

8 Cameron E. A. & Sperandio V. Frenemies: Signaling and nutritional integration in pathogen-microbiota-host interactions. Cell Host & Microbe 18, 275–284 (2015).

9 Rahme L. G. et al. Common virulence factors for bacterial pathogenicity in plants and animals. Science 268, 1899–1902 (1995).

10 Barth A. L. & Pitt T.L. The high amino acid content of sputum from cystic fibrosis patients promotes growth of auxotrophic *Pseudomonas aeruginosa*. J. Med. Microbiol. 45, 110–119 (1996)

11 Field T.R., Sibley C.D., Parkins M.D., Rabin H.R. & Surette M.G. The genus *Prevotella* in cystic fibrosis airways. Anaerobe 16, 337–344 (2010).

12 Tunney M. M. et al. Detection of anaerobic bacteria in high numbers in sputum from patients with cystic fibrosis. Am. J. Resp. Crit. Care Med. 177, 995–1001 (2008).

13 Bassis C. M. et al. Analysis of the upper respiratory tract microbiotas as the source of the lung and gastric microbiotas in healthy individuals. mBio 6, e00037–15 (2015).

14 Goddard A. et al. Direct sampling of cystic fibrosis lungs indicates that DNA-based analyses of upper-airway specimens can misrepresent lung microbiota. Proc. Nat. Acad. Sci. U.S.A. 109, 13769–13774 (2012).

15 Whiteson K. L. et al. The upper respiratory tract as a microbial source for pulmonary infections in cystic fibrosis. Parallels from island biogeography. Am. J. Resp. Crit. Care Med. 189, 1309–1315 (2014).

16 Bradshaw D.J., Homer K.A., Marsh P.D. & Beighton D. Metabolic cooperation in oral microbial communities during growth on mucin. Microbiology 140, 3407–3412 (1994).

17 Flint H. J. et al. Microbial degradation of complex carbohydrates in the gut. Gut Microbes 3, 289–306 (2012).

18 Garrity G., Brenner D.J., Krieg N.R. & Stanley J.R. Bergey's Manual of Systematic Bacteriology. Vol 2: the proteobacteria, Part B: the gammaproteobacteria (Springer, 2005).

19 Staudinger B. J. et al. Conditions associated with the cystic fibrosis defect promote chronic *Pseudomonas aeruginosa* infection. Am. J. Resp. Crit. Care Med. 189, 812–824 (2014).

20 Jorth P. et al. Regional isolation drives bacterial diversification within cystic fibrosis lungs. Cell Host & Microb 18, 307–319 (2015).

21 Willner D. et al. Spatial distribution of microbial communities in the cystic fibrosis lung. ISME J. 6, 471–474 (2012).

22 Palmer K.L., Mashburn L.M., Singh P.K. & M Whiteley. Cystic fibrosis sputum supports growth and cues key aspects of *P. aeruginosa* physiology. J. Bacteriol. 187, 5267–5277 (2005).

23 Niederman R. et al. Short-chain carboxylic acid concentration in human gingival crevicular fluid. J. Dental Res. 76, 575–579 (1997).

24 Korgaonkar A. K. & Whiteley M. Pseudomonas aeruginosa enhances production of antimicrobial in response to N-acetylglucosamine and peptidoglycan. J. Bacteriol. 193, 909–917 (2011).

25 LaBauve A. E. & M.J. Wargo Detection of host-derived sphingosine by *Pseudomonas aeruginosa* is important for survival in the murine lung. PLoS Pathogens 10, e1003889 (2014).

26 Aristoteli L. & Wilcox M.D.P. Mucin degradation mechanisms by distinct *Pseudomonas aeruginosa* isolates. Infect. Immun. 71, 5565–5575 (2003).

27 Robinson C. V. et al. Desulfurization of mucin by *Pseudomonas aeruginosa*: influence of sulfate in the lungs of cystic fibrosis patients. J. Med. Microbiol. 61, 1644–1653 (2012).

28 Ng K. M. et al. Microbiota-liberated host sugars facilitate post-antibiotic expansion of enteric pathogens. Nature 502, 96–99 (2013).

29 Fourrier F., Duvivier B., Boutigny H., Roussel-Delvallez M. & Chopin C. Colonization of dental plaque: a source of nosocomial infections in intensive care unit patients. Crit. Care Med. 26, 301–308 (1998).

30 Scannapieco F.A., Stewart E.M. & Mylotte J.M. Colonization of dental plaque by respiratory pathogens in medical intensive care patients. Crit. Care Med. 20, 740–745 (1992)

31 Scannapieco F. & Mylotte J.M. Relationships between periodontal diseases in the institutionalized elderly with a special focus on pneumonia. J. Periodontol. 67, 1114–1122 (1996).

32 Ghorbani P. et al. Short chain fatty acids affect cystic fibrosis airway inflammation and bacterial growth. Eur. Respir. J. ERJ–01436 (2015).

33 Mirkovic B. et al. The role of short-chain fatty acids, produced by anaerobic bacteria, in the cystic fibrosis airway. Am. J. Resp. Crit. Care Med. 192, 1314–1324 (2015).

34 Høverstad T. & Midtvedt T. Short-chain fatty acids in germfree mice and rats. J. Nutr. 116, 1772–1776 (1986).

35 Hampton T. H. et al. The microbiome in pediatric cystic fibrosis patients: the role of shared environment suggests a window of intervention. Microbiome 2, 14 (2014).

36 Madan J. C. et al. Serial analysis of the gut and respiratory microbiome in cystic fibrosis in infancy: Interaction between intestinal and respiratory tracts and impact of nutritional exposures. mBio 3, e00251–12 (2012).

37 Ramsey B. W. et al. Intermittent administration of inhaled tobramycin in patients with cystic fibrosis. N. Engl. J. Med. 340, 23–30 (1999).

38 Saiman L. et al. Effect of azithromycin on pulmonary function in patients with cystic fibrosis uninfected with *Pseudomonas aeruginosa*: a randomized controlled trial. JAMA 303, 1707–15 (2010).

39 Hurley M. N. et al. Results of antibiotic susceptibility testing do not influence clinical outcome in children with cystic fibrosis. J. Cyst. Fibros. 11, 288–92 (2012).

40 Klepac-Ceraj Ceraj et al. Relationship between cystic fibrosis respiratory tract bacterial communities and age, genotype, antibiotics, and *Pseudomonas aeruginosa*. Environ. Microbiol. 12, 1293–1303 (2010).

41 McDonald D. M. Angiogenesis and remodeling of airway vasculature in chronic inflammation. Am. J. Respir. Crit. Care Med. 164, 39–45 (2001).

42 Mayer-Hamblett Hamblett et al. Association between pulmonary function and sputum biomarkers in cystic fibrosis. Am. J. Respir. Crit. Care Med. 175, 822–828 (2007).

43 Henke M. O. et al. Serine proteases degrade airway mucins in cystic fibrosis. Infect. Immun. 79, 3438–3444 (2011).

44 Badalamenti J. & Hunter R.C. Complete genome sequence of *Achromobacter xylosoxidans* MN001, a cystic fibrosis airway isolate. Genome Announc. 3, e00947–15 (2015).

45 Marsili E. et al. Microbial Biofilm Voltammetry: Direct Electrochemical Characterization of Catalytic Electrode-Attached Biofilms. Appl. Environ. Microbiol. 74, 7329–37 (2008).

46 Spilker T., Coenye T., Vandamme P. & Lipuma J.J. PCR-based assay for differentiation of *Pseudomonas aeruginosa* from other *Pseudomonas* species recovered from cystic fibrosis patients. J. Clin. Microbiol. 42, 2074–2079 (2004).

47 Caporaso J. G. et al. QIIME allows analysis of high-throughput community sequencing data. Nat. Methods 7, 335–336 (2010).

48 Masella A. P. PANDAseq: paired-end assembler for Illumina sequences. BMC Bioinformatics 13, 31 (2012).

49 DeSantis TZ et al. (2006) Greengenes, a chimera-checked 16S rRNA gene database and workbench compatible with ARB. Appl Environ Microbiol 72:5069–5072.

50 White J.R., Nagarajan N. & Pop M. Statistical methods for detecting differentially abundant features in clinical metagenomic samples. PLoS Comput. Biol. 5, e1000352 (2009).

51 Hunter R. C. et al. Ferrous iron is a significant component of bioavailable iron in the cystic fibrosis airways. mBio 4, e00557–13. (2013)

